# Qing-Luo-Yin-induced SIRT1 inhibition contributes to the immune improvement of adjuvant-induced arthritis rats

**DOI:** 10.1101/2024.09.13.608378

**Authors:** Dan-Dan Wang, Meng-Ke Song, Qin Yin, Wen-Gang Chen, Opeyemi Joshua Olatunji, Kui Yang, Jian Zuo

## Abstract

The herbal formula Qing-Luo-Yin (QLY) was proved containing SIRT1 inhibitors. Whether they contribute to the anti-rheumatic effects is to be confirmed. Adjuvant-induced arthritis (AIA) rats were treated by QLY or/and nicotinamide mononucleotide (NMN) for 38 days. After sacrifice, main tissues were collected for histological and western-blot experiments. Levels of rheumatoid arthritis (RA)-related indictors in blood or tissue homogenates were detected by commercial kits. Normal pre-adipocytes were cultured by the relevant rat serums, and the medium was collected for monocytes culture. In replicate experiments, some pre-adipocytes received additional compounds or SIRT1 silencing/overexpression treatments. Due to spontaneous remission of inflammation, QLY didn’t further improve immune milieu in AIA rats, but greatly eased paw edema and joint injuries. Besides, it reversed triglyceride/glucose depletion in liver and adipose tissues, and inhibited the expression and function of SIRT1, causing concomitant changes of related signals and adipkines production. All the effects were weakened by NMN, which activated SIRT1 by increasing NAD production. The serum from QLY-treated rats improved AIA rat serum-induced metabolism and secretion changes of pre-adipocytes, and reduced the secretion of IL-1β, IL-6 and TNF-α in the monocytes cultured with the corresponding medium. A mixture of matrine, sinomenine, sophocarpine, dioscin, berberine showed the similar effects on pre-adipocytes to QLY-containing serum. eNAMPT decrease was especially notable, which was obviously weakened by SIRT1 overexpression but overshadowed SIRT1-silencing. SIRT1 inhibitors in QLY reshaped metabolism and secretion profiles of adipose tissues. It consequently mitigated eNAMPT-mediated inflammation and eased AIA in rats.

## Introduction

Rheumatoid arthritis (RA) is characterized by an initial state of autoimmune reactions, which lead to persistent inflammation within joints or other affected organs.^1^ Clinical presentations of the disease primarily include synovial invasion and joint injuries. According to the current understanding, RA progression is mainly driven by T cells. Their hyper-activation within synovial tissue finally culminates in the evident tissue degradation and inflammation. These outcomes are the direct results from the disruption of cytokines network. Increase of inflammatory cytokines like IL-1, IL-6, IL-17 and TNF-α is now recognized as the key inducer of RA-related manifestations.^2^ But we should realize that the relevant cytokines are not exclusively produced by T cells. Monocytes/macrophages are another key source. Being an integral component of the innate immune system, they are responsible for the clearance of pathogens and antigen presentation. They extensively interact with lymphocytes, and play multiple roles in RA pathology.^3–4^ Therefore, it is interesting to investigate their changes in RA. Monocytes are deemed as the main source of residential macrophages. Because of the accessibility reason, they are preferentially investigated in related researches.

M1 macrophages act as a contributory factor in inflammation. According to this, the polarization profile of monocytes in RA patients significantly diverges from that of healthy individuals, demonstrating a phenotype resembling to M1 macrophages. There is a marked increase in the frequency of inflammatory CD16^+^ monocytes, and it positively correlates with RA severity.^5^ Monocytes/macrophages are adept at recruiting T cells and fostering their differentiation towards inflammatory phenotypes within RA synovium.^6^ In fact, the interaction is reciprocal. Certain T cell subtypes possess the ability to recruit monocytes/macrophages, promote the secretion of inflammatory cytokines, and induce their differentiation toward osteoblasts.^7^ In this context, inhibiting inflammatory polarization of monocytes/macrophages will disrupt this malicious feedback loop, and effectively alleviate RA.^8–10^

Increasing evidence indicates that beyond traditional immune mediators, energy metabolism status represents another critical determinant governing immune functions of monocytes/macrophages.^11^ Their inflammatory polarization relies on glycolytic pathway.^12^ These clues suggest that metabolism regulation system and the related organs would play a role in reshaping monocytes/macrophages phenotypes. White adipose tissues (WAT) are non-negligible in this aspect. Similar to liver, WAT is a key metabolism hub. Being the fat reservoir, it provides sustainable energy supply for the energy-consuming cells, including monocytes/macrophages, when it is necessary. For example, WAT undergoes lipolysis to release more fatty acids to meet the increased energy requirement of the immune system in RA patients.^13^ Furthermore, WAT would participate in immune regulation by secreting adipokines. These substances are involved in both metabolism and immune regulation. Taking extracellular NAMPT (eNAMPT) an example, it promotes monocytes/macrophages-mediated inflammation by activating TLR4.^14^ Besides, it facilitates glycolysis as a rate-limiting enzyme in synthesizing NAD.^15^ Accordingly, its abnormal increase is deeply implicated in inflammatory diseases, and it has been recognized as a promising therapeutic target of RA.^16^ Hence, the signal transduction mechanism related to eNAMPT should be highlighted.

eNAMPT is the secreted form of intracellular NAMPT (iNAMPT). It affects the internal environment basically by providing NAD to activate SIRT1, a deacetylase. It has been confirmed that SIRT1 serves as the most important energy sensor, regulating PPARγ, PGC-1α and some other factors to favor oxidative phosphorylation. Via deacetylation, SIRT1 inhibits the activities of many inflammatory transcriptional factors, such as AP-1, HIF-1α and NF-κB.^17^ Because of these properties, SIRT1 activation is usually believed to be beneficial for anti-RA therapies.^18^ However, convincing evidence show that SIRT1 overexpression in RA patients’ synovial tissues is common, where SIRT1 induces inflammatory secretion and apoptosis resistance.^19^ Our previous studies also show that SIRT1 up-regulation in peripheral immune cells is a sign of acute inflammation during active RA.^20–21^ These facts reveal the other facet of SIRT1. By taking the inflammatory nature of its upstream NAMPT into consideration, we can conclude that NAMPT/SIRT1 has dual impacts on the immune system. A proof is that SIRT1 accounts for eNAMPT processing and production.^22^ Via this manner, SIRT1 would indirectly provoke inflammatory reactions.

The above clues hint that SIRT1 inhibition may be conditionally favorable for RA patients, especially in WAT. Unfortunately, almost all the related researches focus on SIRT1 activation, thanks to the well-acknowledged anti-inflammatory outcomes. To fully clarify the role of NAMPT/SIRT1 in RA and exploit the clinical application, it is necessary to investigate the impacts of SIRT1 inhibitors during the therapies. We previously confirmed that activating SIRT1 can indeed attenuate the severity of RA animal models.^23–24^ Interestingly, our recent work has showed that a Traditional Chinese Medicine (TCM) anti-rheumatic formula Qing-Luo-Yin (QLY) contains SIRT1 inhibitors.^25–27^ Traditional Chinese medicine(TCM) has unique advantages in the prevention and treatment of diseases owing to its holistic view and more than 2 000 years of experience in the clinical use of natural medicine.^28–29^ This formula was developed by a contemporary TCM master Jiren Li, and demonstrates impressive effects in treating hot syndromes-related RA.^30^ QLY profoundly affect metabolism and immune conditions in rheumatic subjects, and WAT is one of its targets.^29–30^ Given the pivotal role of SIRT1 in determining internal environment, the SIRT1 inhibitors in QLY must be related to the observed outcomes. This study was designed to test this assumption using an adjuvant-induced arthritis (AIA) model. Obtained results confirm that QLY-induced SIRT1 inhibition in AIA rats’ WAT contributes to the therapeutic effects by reducing eNAMPT production and then affecting the phenotype of monocytes.

## Material and methods

### Reagents

Heat-inactivated Bacillus Calmette-Guérin (BCG) and incomplete Freund’s adjuvant (IFA) were supplied by Rebio Scientific (Shanghai, China). ELISA quantification kits of IL-1β, IL-6, IL-17A, MCP-1, TNF-α, leptin, and adiponectin were purchased from MultiScience Biotech (Hangzhou, Zhejiang, China). Colorimetric quantification kits of reduced glutathione (GSH), malonaldehyde (MDA), superoxide dismutase (SOD), alkaline phosphatase (AKP), glycated serum protein (GSP), glucose (GLU), lactic acid, triglyceride (TG), total cholesterol (T-CHO), high-density lipoprotein cholesterol (HDL-C), and low-density lipoprotein cholesterol (LDL-C) together with eNAMPT ELSIA kit were supplied by Jiancheng Bioengineering Institute (Nanjing, Jiangsu, China). SIRT1 overexpression plasmid was constructed by General Biol (Chuzhou, Anhui, China) based on pcDNA3.1-EGFP vector. The primers and siRNA used were synthesized by Universal Biology (Chuzhou, Anhui, China), and their sequences are shown in **Supplementary S1**. Nicotinamide mononucleotide (NMN) was procured from Sigma-Aldrich (St Louis, MO, USA). QLY-related compounds matrine, sinomenine, sophocarpine, dioscin, berberine were the products of Bencao Yikang Biotechnology (Nanjing, Jiangsu, China). The other reagents used were all from domestic suppliers.

### Preparation and administration of QLY extract

Raw herbal drugs including Kushen (radix of *Sophora flavescens* Ait.), Qingfengteng [caulis of *Sinomenium acutum* (Thunb.) Rehder and E. H. Wilson], Huangbai (cortex of *Phellodendron chinense* C. K. Schneid.), and Bixie [rhizome of *Dioscorea collettii var. hypoglauca* (Palib.) S. J. Pei and C. T. Ting] were brought from Bozhou Herbal Medicine Market (Anhui, China), and authenticated by Professor Jian Zuo (Wannan Medical College, Anhui, China). Their voucher specimens were kept in the Herbarium Center of Wannan Medical College (ID: 2022-013). These ingredients were combined in a ratio of 1.5: 1.2: 1: 1, and pulverized. The powder underwent reflux extraction with 95% ethanol twice. The remaining residues were further boiled in water for one time. The combined filtrates were dried to yield a sticky QLY extract.

For the administration use, this product was dissolved in ethanol, and then evenly dispersed in 0.5% CMC-Na solution. Ethanol content in the suspension is about 1.2%. Animals in the following experiment were orally administered with this suspension. For the sake of animal welfare, we minimized rats use in this study, and only one dose was adopted, because the pharmacological properties of QLY had been well studied. According to body surface area-based equivalent dose calculation and the previous studies, we used an optimal dose (equivalent to 0.45 g/kg of raw drugs).^14,29^

### Establishment of AIA model in rats

The animal experiment protocol received the approval from the Ethics Committee of Wannan Medical College (Ethics Approval Number: WNMC-AWE-2023358). All the experimental procedures were strictly adhered to ARRIVE guidelines. A group of 32 male SD rats (7-weeks old, obtained from Tianqin Biotechnology Co., Ltd.) were housed in a specific-pathogens free laboratory, and acclimated for 1 week. BCG was first grinded in a mortar carefully, and mixed with IFA to prepare complete Freund’s adjuvant (CFA, with BCG concentration of 10mg/ml). Twenty-four rats then received the intradermal injection of CFA (0.1 ml) into the plantar skin of the left hind paw. They were assigned into three groups: AIA models (AIA), QLY-treated AIA rats (QLY), and QLY plus NMN-treated AIA rats (QLY+NMN). The untreated 8 rats were adopted as normal healthy controls (normal). All the treatments were administered by gavage, and the normal and AIA controls were treated by 0.5% NMC-Na. Based on the previous study, the dose of NMN was set at 200 mg/kg.^31^

After the continuous treatments for 38 days, the maximum amounts of rat blood were collected into anticoagulant and coagulation-promoting tubes through abdominal aorta until euthanasia under pentobarbital sodium-induced anesthesia status. After centrifugations, plasma and serum were obtained, which were used for the following tests or cell culture. The hind paws, liver, and perirenal fat pads were removed, and weighed. Parts of livers were subjected to Oil Red O staining-based histological examination. The other tissue samples were dyed with hematoxylin-eosin (H&E) dyes. The experiments followed the same steps as previously described.^32–33^ Possible abnormalities of tissue and cell morphology were observed by the aid of an Olympus BH-2 optical microscope (Tokyo, Japan). Expression of SIRT1, PPARγ, PGC-1α, HSL and IL-1β in WAT homogenates were investigated by western-blot (WB) method. Ace-PPARγ was detected by immunoprecipitation (IP) method. RA-related indicators in blood or liver homogenates were determined by commercial kits in accordance to the manufacturers’ instructions.

### Isolation and culture of rat primary monocytes and preadipocyte

The unicellular suspension resulted from collagenase I-based WAT digestion was first centrifuged at 500 g for 10 min. The precipitated cells were re-suspended and cultured in DMEM/F12 medium supplemented with 10% FBS and penicillin-streptomycin (100 U/mL) at 37°C in a humidified atmosphere with 5% CO_2_. When reaching 80% of confluence, the cells were passaged in a ratio of 1: 3. Homogeneous fibroblast-like pre-adipocytes from the 3^rd^ to 8^th^ passages were used in the following experiments.

Monocytes were isolated from the fresh rat blood using a separation kit provided by Solarbio Biotech (Beijing, China). By the aid of the provided reagents in the kit, the centrifugation at 500 g for 20 min generated a cellular layer, which was carefully aspirated and washed twice. The cells were then cultured in the normal circumstance, and the attached ones were taken as monocytes. Expression levels of genes involved in their polarization were assessed by reverse transcription polymerase chain reaction (RT-PCR) assays. The remaining cells were used in the experiments in vitro. All these cells were used for only one time without further passage.

### Treatments of cells in vitro

Pre-adipocytes were seeded in 24-well plates under a sterile condition. The initial complete medium used was comprised of 10% FBS. After the cells adhered to wall, the medium was replaced by the fresh one containing 10% different serums obtained from normal healthy, AIA model and QLY-treated AIA rats. Twenty-four hours later, the pre-adipocytes were collected for WB experiments, while the levels of certain metabolites and cytokines in the medium were determined by the corresponding kits. The mediums were then used to culture monocytes from healthy rats for 24 h. The medium from this experiment was also collected for ELSIA analysis. In the replicate experiment, we only cultured rat pre-adipocytes with different serums, and performed immunofluorescence assay after the culture.

In the next experiment, pre-adipocytes in 6-wells plates were first cultured in the medium supplemented by either normal or AIA rat serum for 12 h. Then, half of the medium was replaced by the fresh ones comprised of normal rats’ serum, AIA rats’ serum, AIA rats’ serum + QLY-related compounds or QLY-treated AIA rats’ serum. After the further 24 h culture, the cells were collected for WB assays, and the medium was collected for biochemical or ELISA analyses. A mixture comprised of matrine (100 ng/ml), sinomenine (80 ng/ml), sophocarpine (100 ng/ml), dioscin (25 μg/ml), and berberine (25 μg/ml) was used in the compounds treatment.

To clarify the role of SIRT1 in QLY-caused changes of pre-adipocytes, this gene was silenced or overexpressed prior to the treatments. The cells were seeded in 6-well plates. When reaching 40% of confluence, some cells were treated by siRNA-SIRT1 (100 nM) along with Lipofectamine 2000 reagent (Invitrogen, Shanghai, China). In another experiment, SIRT1 overexpression plasmid (100 nM) was transfected into the cells by using Lipofectamine 3000 reagent (Invitrogen, Shanghai, China). The other arrangements were the same with those depicted above.

### RT-PCR assay

This assay was conducted based on the previous reported protocols.^34–35^ Trizol reagent (Tiangen, Beijing, China) was used to extract total RNA from cells. RNA was reverse-transcribed into complementary DNA (cDNA) using a synthesis kit. For the quantitative PCR (qPCR) purpose, Universal qPCR Master Mix kit was used, and the experiment was conducted on a 7500 Real-Time PCR Detection system (Applied Biosystems, Life Technologies). The synthesized cDNA served as templates, and the relative mRNA expression levels were calculated by using the 2^^(-ΔΔCT)^ method, with *β-Actin* as the internal reference gene.

### WB and IP assays

Tissue or cell samples were added into RIPA buffer containing PMSF and a protease inhibitor mixture (Beyotime, Shanghai, China). The mixture was then lysed on ice for 30 min to 2 h, and centrifuged to extract the supernatant. Protein concentrations in the supernatants were determined by a BCA assay kit (Beyotime, Shanghai, China). The protein samples underwent electrophoresis on an 8-10% SDS polyacrylamide gel, and the separated proteins were transferred onto nitrocellulose membranes (NC, Millipore, USA) in ice-water. After being blocked with 5% bovine serum albumin (BSA), the membranes were incubated overnight at 4°C with primary antibodies. Afterwards, they were washed three times, followed by incubation with appropriate HRP-conjugated IgG secondary antibodies at room temperature for 1 h.^36^ The protein signals were detected using the Amersham Imager 600 ECL Imaging System. Relative expression levels of protein were normalized to β-Actin or GAPDH by the aid of Image J software (version 1.52a, NIH, Bethesda, MD, USA).

In IP assay, some of the above lysates were incubated protein A/G magnetic beads in the diluted primary acetylated pan antibody solution at 4°C overnight. The proteins bound to the beads were thoroughly extracted by boiling protein loading buffer. PPARγ concentrations in this product were assessed by WB method.^37–38^ All the raw images from these experiments are shown in **Supplementary S2.**

### Immunofluorescence Experiment

By the end of cell culture in an experiment above, pre-adipocytes were first treated by 0.1% Triton solution for 10 min. After extensive washing, 5% BSA-based blocking procedure was performed. Next, primary rabbit anti-rat NAMPT and mouse anti-rat acetylated pan antibodies (1: 100) were added. An overnight incubation at 4°C was carried out. The cell samples were then treated with fluorescein-labeled secondary IgG antibodies (1: 500) at room temperature for 1 h. Finally, the cell nuclei were stained with DAPI. After being sealed with the anti-fade mounting reagent, the cells were observed by a SP8 Lightning confocal microscope. In the replicate experiment, the cells were incubated in primary rabbit anti-rat IL-1β and mouse anti-rat PPARγ antibodies (1: 100) after BSA blocking.

### Statistical Analysis

Mean values along with their corresponding standard deviations were used to present the data. Statistical comparison among groups was performed using one-way analysis of variance (ANOVA) followed by Tukey’s post hoc test by the aid of GraphPad Prism 8.0 (GraphPad Software, Cary, NC, USA).

## Results

### SIRT1 inhibition was involved in the therapeutic actions of QLY on AIA rats

SIRT1 activity relies on NAD, but cells usually can’t utilize extracellular NAD. In this study, we used NMN, the synthetic precursor of NAD to promote SIRT1 functions in some QLY-treated AIA rats. By the end of the in vivo experiment, inflammation had been spontaneously eased a lot. There was no obvious change about the expression of gene *IL-1β*, *iNOS*, *IL-10*, *Arg-1*, *MCP-1*, *SIRT1* and *PPARγ* in circulating monocytes of AIA rats, compared with the normal controls. Interestingly, *IL-6* expression was even decreased. Under this context, neither QLY nor QLY+NMN treatments affected the expression of these genes, including *SIRT1* (**Figure 1A**). The similar results were observed about interleukins in plasma. The concentrations of IL-1β, IL-6, and IL-17A in the all the rats’ plasma were similar. However, AIA-induced MCP-1 increase was persistent, which was reduced by QLY (**Figure 1B**). NMN showed no impact on the result, suggesting the weak effects of SIRT1 on MCP-1 production. Oxidative stress markers GSH, MDA, and SOD also showed no significant differences among the groups (**Figure 1C**). Despite the spontaneous remission of AIA-induced inflammation, hind paw edema was still evident. QLY significantly reduced the weight index of rat hind paws in AIA rats, and NMN reversed this effect (**Figure 1D**). Levels differences of the bone remodeling indicator AKP among these groups were generally similar to paw edema. Although not significant, NMN offset QLY-induced AKP decrease in AIA rats too (**Figure 1E**). Examination of H&E stained-sections reveals that AIA rats suffered from obvious cartilage erosion and synovial hyperplasia. QLY treatment alleviated these abnormalities, but cartilage degradation can be still observed in the AIA rats receiving QLY+NMN treatment (Figure 1F). The results preliminarily show the relevance of SIRT1 inhibition with QLY-induced therapeutic effects on AIA rats, and suggest that immune cells like monocytes may not the preferential targets of QLY.

**Figure 1.**
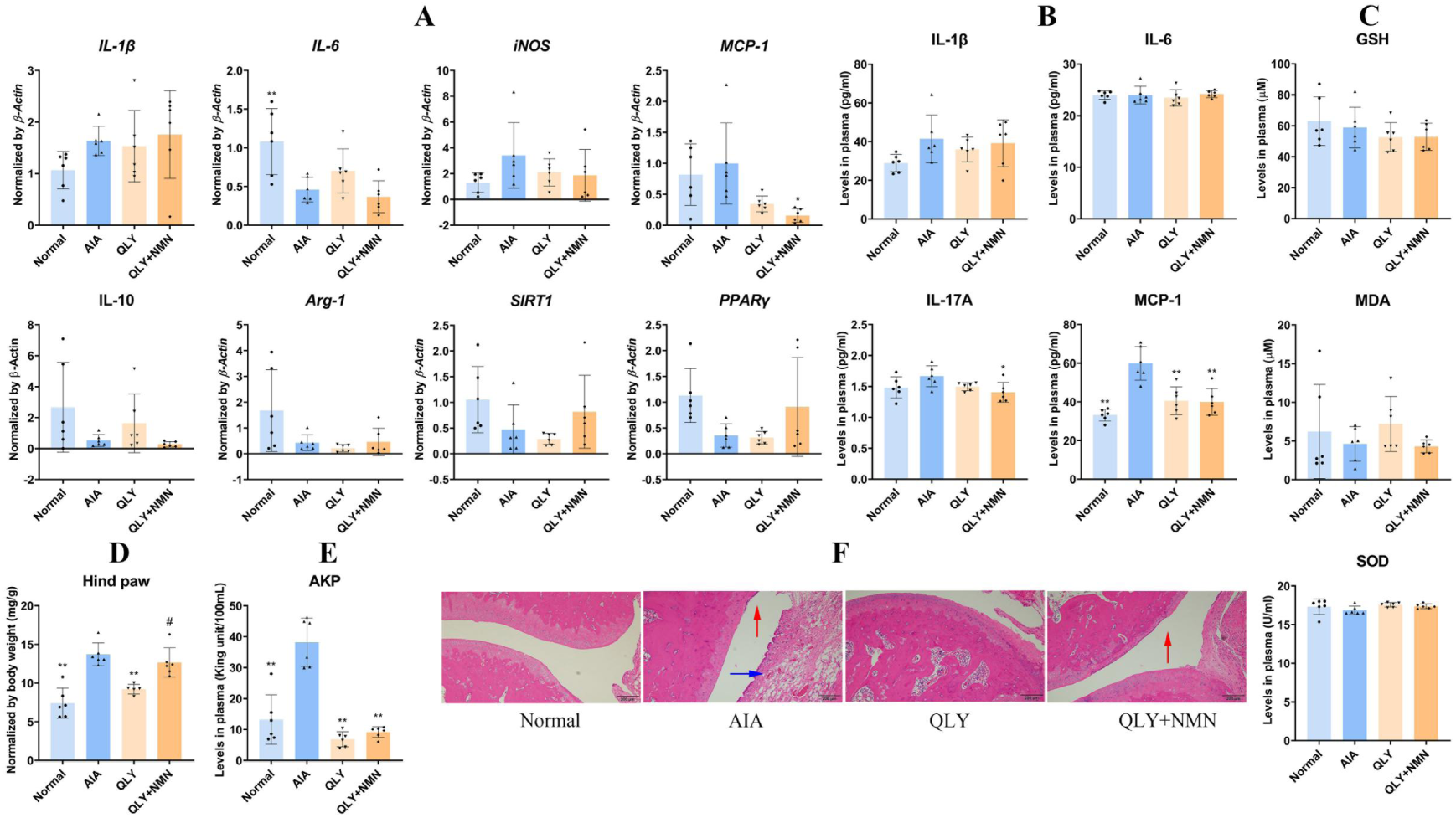
QLY therapy improved arthritic manifestations of AIA rats. A, expression of some polarization-related genes in blood monocytes; B, levels of some RA-related cytokines (IL-1β, IL-6, IL-17A and MCP-1) in the rats’ blood; C, levels of oxidative indicators (GSH, MDA and SOD) in the rats’ blood; D, the relative weight of left hind paw; E, AKP levels in the rats’ blood; F, H&E staining-based histological examination of the left hind ankle joint (the red and blue arrows indicate cartilage erosion and synovial invasion, respectively). Statistical significance: *p < 0.05 and **p < 0.01 compared with AIA model rats; ^#^p < 0.05 compared with QLY-treated AIA rats.

### QLY inhibited SIRT1 in AIA rats’ metabolic organs

SIRT1 is an important metabolic regulator. To confirm the effects of QLY on SIRT1, we detected some metabolites involved in glucose and lipids metabolism. Levels of GSP, GLU and lactic acid in AIA rats’ blood were increased obviously, showing the accelerated glycolysis. Despite TG remained unaffected, levels of T-CHO, HDL-C, and LDL-C were all declined under this condition. We found that QLY generally restored all these changes, and these outcomes were weakened by NMN (**Figure 2A**).

**Figure 2.**
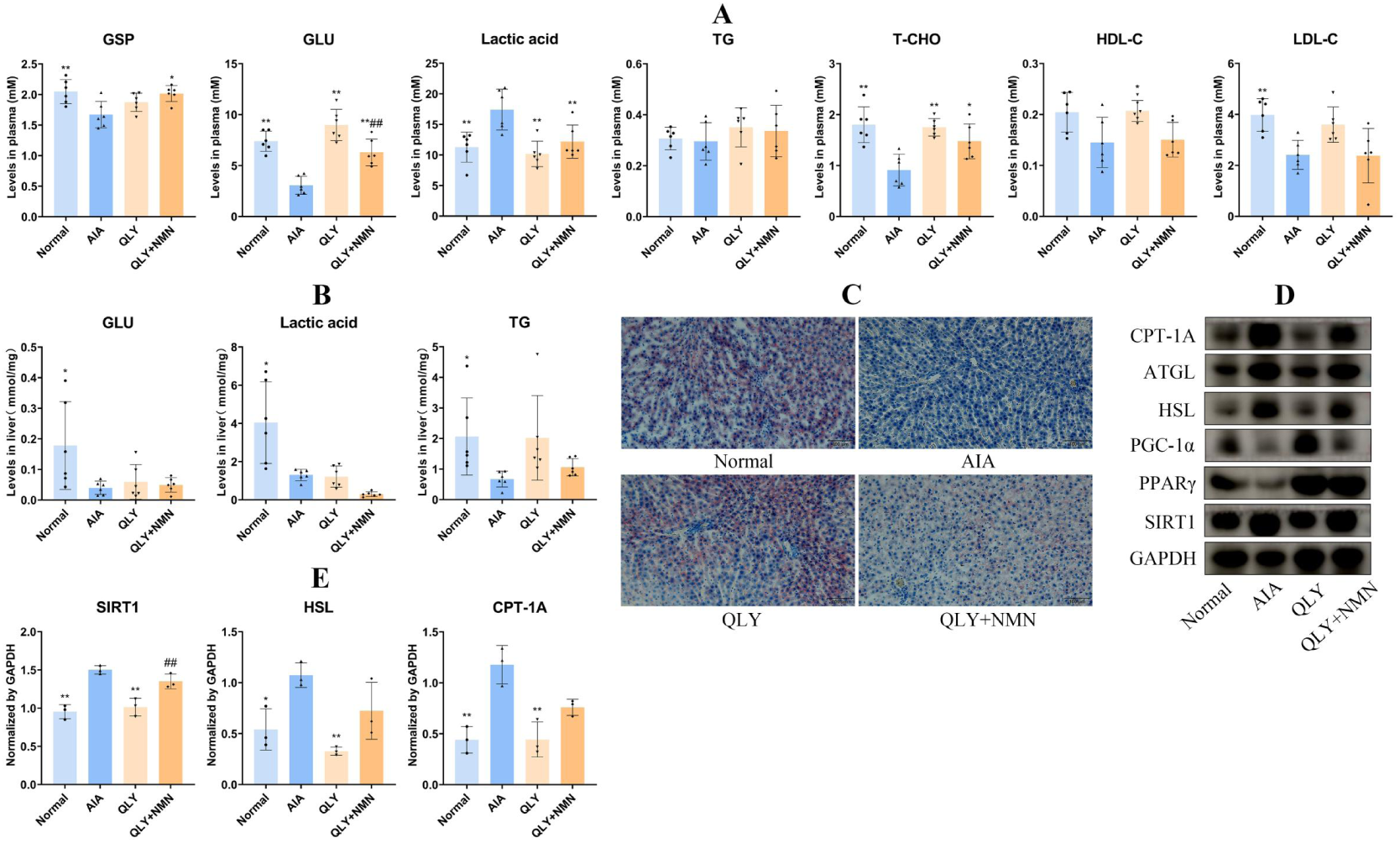
QLY therapy affected SIRT1-related signals as well as metabolic status in AIA rats’ liver. A, levels of glucose and lipids metabolism-related indicators in the rats’ blood; B, levels of some key metabolites (GLU, TG and lactic acid) in the rats’ livers; C, Oil Red O staining-based histological examination of livers; D, WB assay performed using the liver samples; E, the quantification results of assay D. Statistical significance: *p < 0.05 and **p < 0.01 compared with AIA model rats; ^##^p < 0.01 compared with QLY-treated AIA rats.

The above changes are largely determined by liver, the crucial metabolism hub. Hence, we focused on changes there. We first detected some key metabolites related to glucose and lipid metabolism. Liver apparently shouldn’t account for the altered glucose metabolism profile in the rats’ blood, because the changes there were different. Of note, TG was reduced a lot in AIA rats’ liver. This situation was improved by QLY therapy, but this beneficial outcome was impaired by NMN (**Figure 2B**). In line with these results, Oil Red O staining-based histological examination indicates that QLY can significantly restore fat storage decline in AIA rats’ livers, and NMN significantly reversed this effect (**Figure 2C**). Expression changes of SIRT1 and the related signals were much more significant in liver than those observed in blood (**Figure 2D**). SIRT1 as well as fat utilization-related proteins including HSL, ATGL and CPT-1 were overexpressed in AIA rats’ liver, while the expression of PGC-1α and PPARγ was impaired. QLY therapy led to the significant restoration of these changes, which may the reason for the increased TG deposits. NMN antagonized the effects of QLY in this aspect once again (**Figure 2E**). Of note, the above findings reveal that AIA causes different changes on SIRT1 expression in immune cells and liver.

SIRT1 overexpression is one important but not the exclusive factor driving fat catabolism in AIA rats’ liver, and WAT is another key role determining lipids profile. Indeed, fat deposits differences among the rats’ WAT were very significant (**Figure 3A**). As expected, QLY significantly enlarged the volume of shrunk WAT in AIA rats, while NMN exhibited an inhibitory effect against QLY treatment (**Figure 3B**). WAT weight differences must be caused by the changes of adipocytes sized. Adipocytes in AIA rats’ WAT showed the reduced size, and QLY exerted notable therapeutic effects in this regard. NMN counteracted the favorable outcomes brought by QLY (**Figure 3C**). In addition to related metabolic regulators, we detected IL-1β in the WAT-based WB experiments to investigate immune implications of SIRT1 changes. Furthermore, we detected acetylated pan antibody-bound PPARγ using the samples (**Figure 3D**). The expression differences of SIRT1, HSL and other metabolic regulators among the groups were very similar to those observed in liver. Importantly, AIA-caused IL-1β increase was reduced by QLY, which was reversed by NMN (**Figure 3E**). Meanwhile, we noticed that acetylation of PPARγ was facilitated in AIA rats, and QLY hampered this process. The anti-acetylation functions of QLY were thoroughly abrogated by NMN (**Figure 3F**). These facts solidly support that claim that QLY acted as a SIRT1 inhibitor in AIA rats’ WAT. Because WAT is a potent secretion organ, we examined the levels of some adipokines in rat plasma. We found that the levels of adiponectin and leptin were significantly reduced in the plasma of AIA rats, while eNAMPT levels were increased a lot. QLY reversed all the above changes, and its effects on eNAMPT were especially efficient. The combined use of NMN constrained QLY-brought effects to certain extends concerning these parameters (**Figure 3G**). Besides metabolic clues, the changes of ace-PPARγ and eNAMPT confirm that QLY functionally inhibited SIRT1. Considering the secretion capacity of WAT and the immune regulatory role of SIRT1, it would bring profound immune consequences.

**Figure 3.**
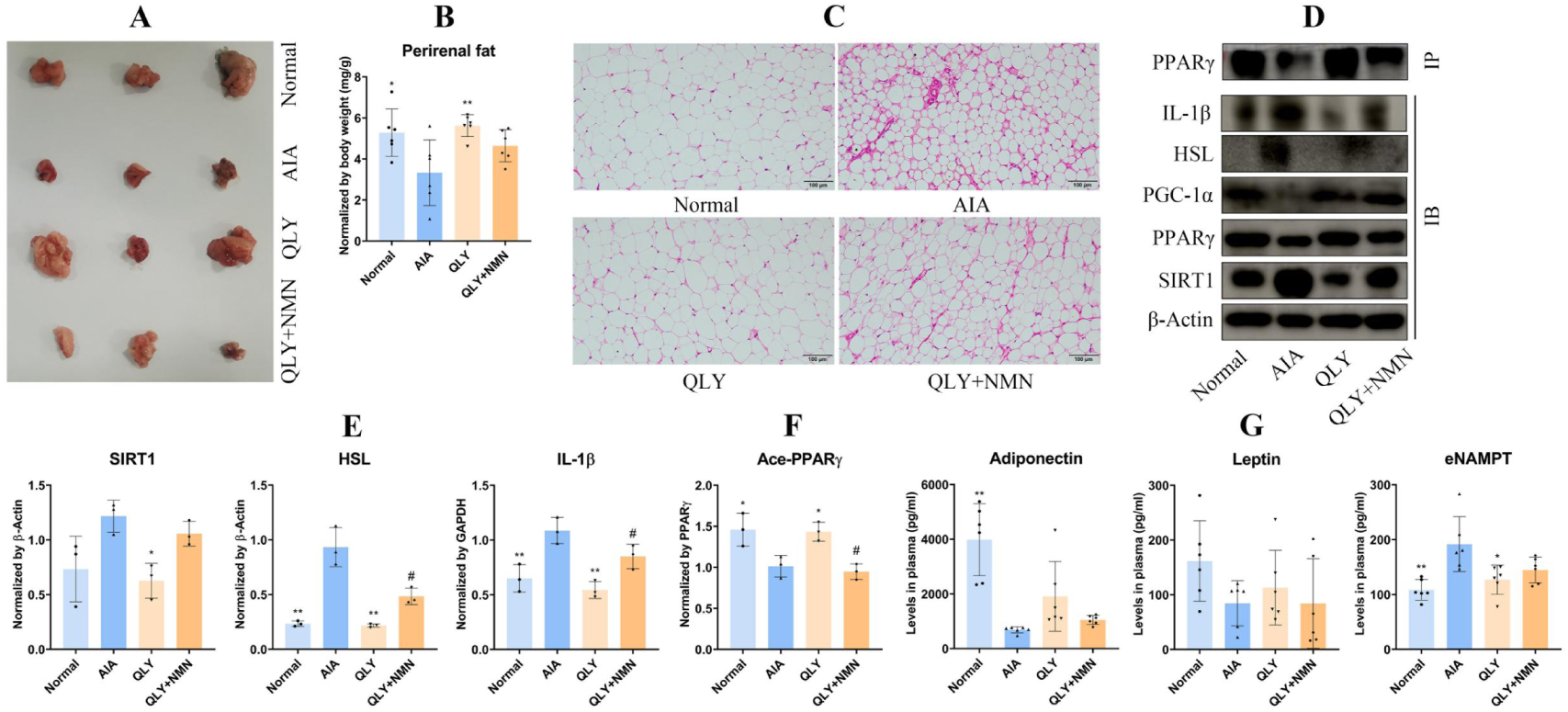
QLY therapy reshaped both metabolism and secretion profiles of WAT in AIA rats by regulating SIRT1 and the related signals. A, morphological observation of the representative perirenal fat pads; B, relative weight of perirenal fat pads; C, H&E staining-based histological examination of perirenal fat pads; D, WB and IP assays performed using the WAT samples; E, the quantification results of WB assays; F, the quantification results of IP assay; G, levels of the representative adipokines (leptin, adiponectin, and eNAMPT) in the rats’ blood. Statistical significance: *p < 0.05 and **p < 0.01 compared with AIA model rats; ^#^p < 0.05 compared with QLY-treated AIA rats.

### QLY eased AIA-induced inflammation in monocytes by reshaping adipocytes

To test the above theory, we isolated monocytes from rat blood and cultured them by using pre-adipocytes-related medium. We first validated the impacts of the different microenvironments on pre-adipocytes using WB method (**Figure 4A**). Similar to the phenomena observed in vivo, AIA condition-caused even more dramatic increase of SIRT1 and HSL. SIRT1 controls the production of eNAMPT, a key inflammatory adipokine. We therefore also investigated total NAMPT (tNAMPT) protein here, whose expression was greatly up-regulated too. QLY suppressed the expression of all the above proteins (**Figure 4B**). Comparatively, the expression differences of PGC-1α and PPARγ were less obvious. The changes led to the varied glycolysis and TG catabolism status. The levels of GLU and TG in the medium from QLY-treated AIA rats’ serum-cultured pre-adipocytes were significantly up-regulated compared with the AIA group, while lactate concentrations were significantly reduced (**Figure 4C**). It further indicates the inhibitory effects of QLY on lipolysis and fat utilization. As the effects of QLY on IL-1β secretion had been confirmed, we herein only detected IL-6 and TNF-α in the medium. Because AIA serum was sampled when inflammation was not intense, it induced only mild increase of IL-6 and TNF-α in the pre-adipocytes. QLY-containing serum significantly decreased their concentrations, validating the anti-inflammatory effect in WAT (**Figure 4D**). Similarly, monocytes released more IL-1β, IL-6, and TNF-α than the controls in the AIA group when co-cultured with pre-adipocytes. The treatment with QLY-containing serum reduced all their increase in the monocyte culture medium (**Figure 4E**). It is noteworthy that the levels variation about these cytokines was much more significant than that observed in the above experiment, showing WAT and immune system-mediated amplification mechanism. It must be mediated by adipokines like eNAMPT.

**Figure 4.**
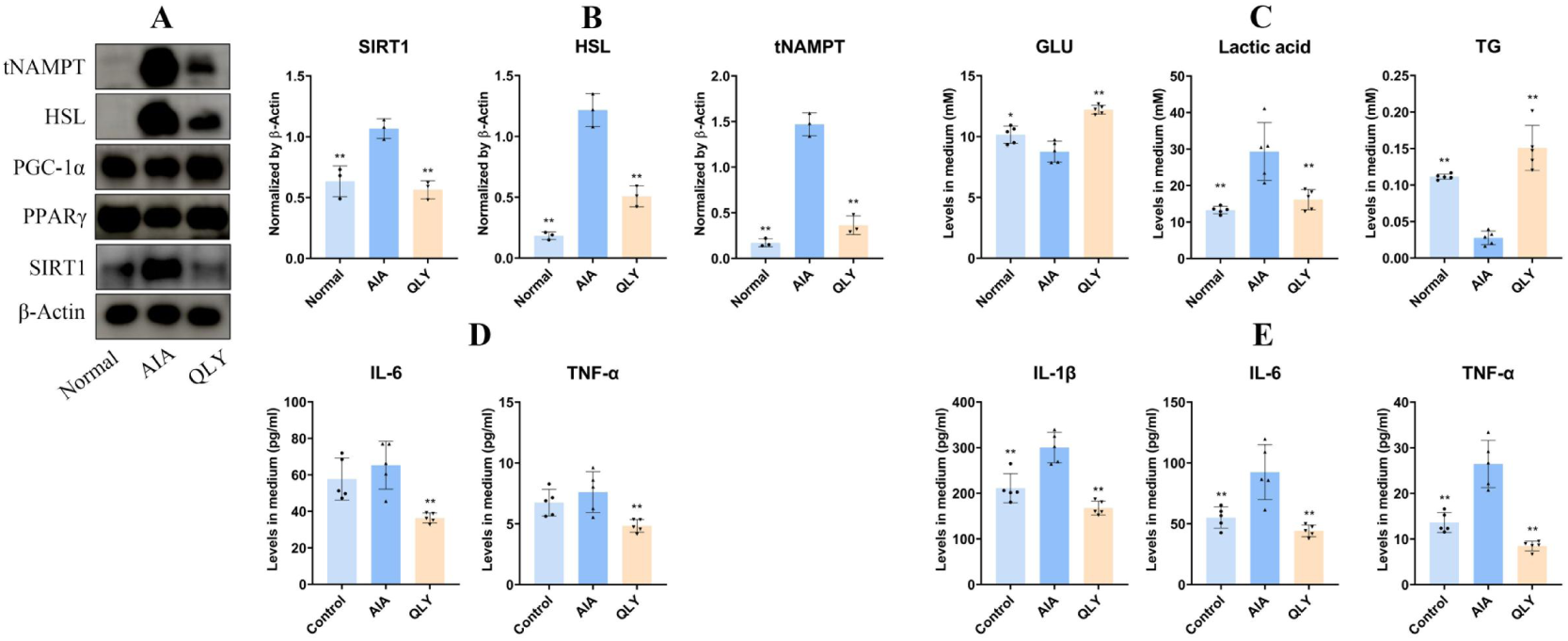
QLY-containing serum-induced changes of pre-adipocytes affected immune status of monocytes. A, WB assays performed using the pre-adipocytes cultured in vitro (they were cultured with the serums from healthy, AIA model or QLY-treated AIA rats); B, the quantification results of assay A; C, levels of some key metabolites (GLU, TG and lactic acid) in the medium from assay A; D, levels of IL-6 and TNF-α in the medium from assay A; E, levels of IL-1β, IL-6 and TNF-α secreted by the monocytes cultured by the medium from assay A. Statistical significance: *p < 0.05 and **p < 0.01 compared with AIA group.

WAT impacts immune environment at least from two approaches, adipokines and cytokines. Although all of they are secreted proteins, their productions are controlled by different mechanisms. We further clarify the mechanisms by which QLY controls the secretion of WAT based on immunofluorescence method. The above results show QLY can potently reduce NAMPT expression. But in fact, NAMPT is dispensable for cell metabolism. The tNAMPT changes should be mainly attributed to eNAMPT other than iNAMPT. Indeed, we observed that iNAMPT expression in pre-adipocytes was largely unaffected by the different rat serums. The varied acetylation status explained the observed tNAMPT expression differences (**Figure 5A**). The overall data show that NAMPT expression in AIA rats is generally up-regulated, but iNAMPT levels remain stable, as more eNAMPT is produced and secreted via SIRT1-mediated acetylation.^20^ IL-1β and many other cytokines are controlled by inflammatory transcriptional factors typically NF-κB. Considering the negative regulation of SIRT1 on NF-κB, it is difficult to explain the inhibitory effects of QLY on these cytokines.^17^ We believed that the favorable outcomes would be mediated by PPARγ, a functional rival of SIRT1 and another anti-inflammatory regulator.^14^ We found that the expression of IL-1β was increased in AIA rat serum-cultured pre-adipocytes, while QLY inhibited this increase a bit. Meanwhile, QLY restored PPARγ expression decline to certain extend and promoted its translocation into nucleus (**Figure 5B**). There, PPARγ will bind to NF-κB and curb its functions.^39^ By taking inhibition of SIRT1 on the expression and functions of PPARγ into consideration, it can be inferred that SIRT1 may be pro-inflammatory in some tissues like WAT, where PPARγ is decisive for the functional phenotypes.

**Figure 5.**
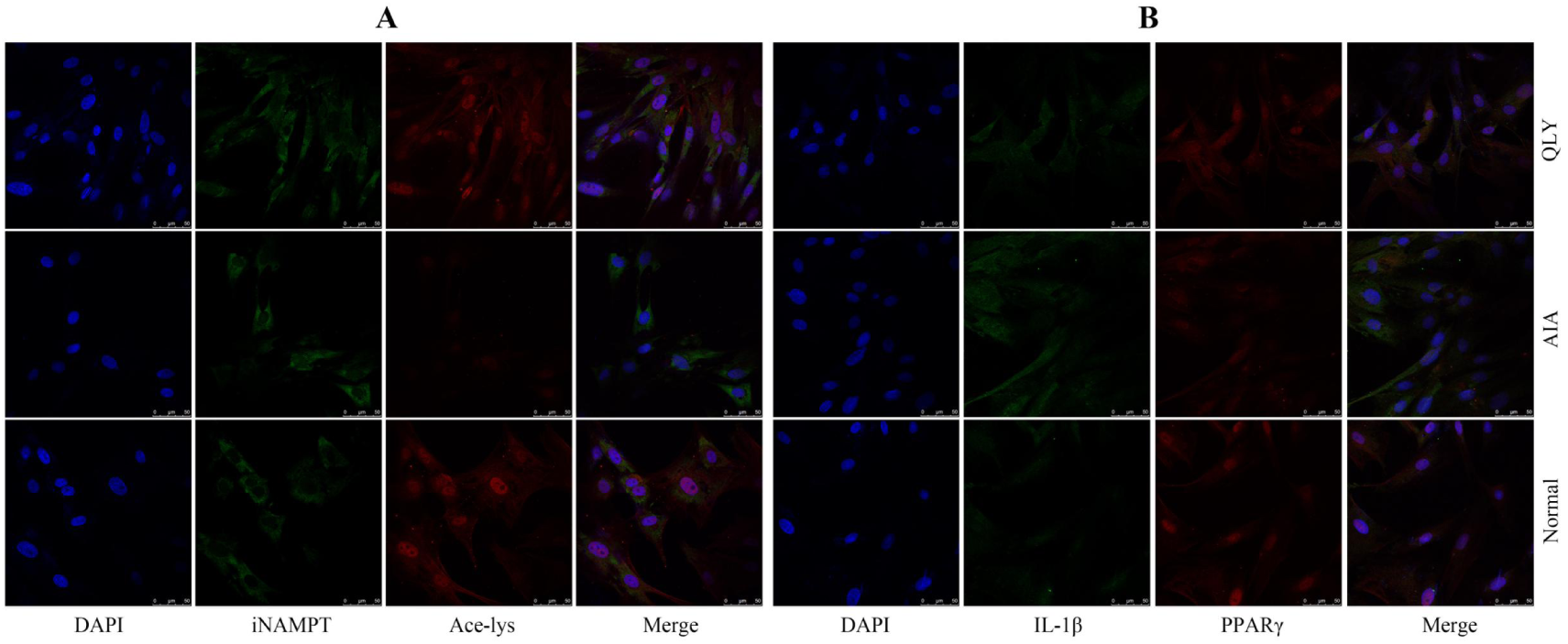
QLY regulated the production of eNAMPT and IL-1β in different manners. A, expression and co-localization of iNAMPT and pan ace-lys in the pre-adipocytes cultured by the serums from normal healthy, AIA model or QLY-treated AIA rats; B, expression and co-localization of IL-1β and PPARγ in the cells receiving the same treatments as above.

### QLY-related compounds prevented inflammatory development of pre-adipocytes by inhibiting SIRT1

Subsequently, we treated pre-adipocytes with QLY-containing serum and some related compounds to confirm whether QLY-induced SIRT1 inhibition indeed prevents the cells from acquiring an inflammatory phenotype. The selection of QLY-derived SIRT1 agonists and their treatment concentrations was based on our previous studies.^10,14,29^ We first investigated their impacts on SIRT1 and its rival PPARγ in the cells in the context of AIA serum stimulus (**Figure 6A**). Similar to QLY-containing serum, the compounds efficiently suppressed SIRT1 protein expression in AIA rat serum-primed pre-adipocytes, but increased the expression of PPARγ as well as PGC-1α (**Figure 6B**). They generally showed the similar impacts on the levels of extracellular GLU, lactic acid and TG too. However, QLY didn’t substantially lactic acid production this time (**Figure 6C**). These results are almost the same as observed in the above assays. It further confirms that SIRT1 inhibitors within QLY suppresses glycolysis and favors TG deposits. In the secretion aspect, both the compounds and QLY-containing serum showed no impacts on adiponectin and leptin. However, they significantly inhibited AIA serum-induced eNAMPT production (**Figure 6D**). This result demonstrates that QLY and the related SIRT1 inhibitors don’t suppress the overall secretion capacity of WAT, but selectively controls eNAMPT release. We also detected some WAT-related abundant cytokines, and revealed that the levels of IL-1β, IL-6 and MCP-1 were all increased by AIA rat serum. QLY and the related compounds generally reduced all the cytokines in this pre-adipocyte culture medium. Comparatively, their effects on IL-1β were weak (**Figure 6E**).

**Figure 6.**
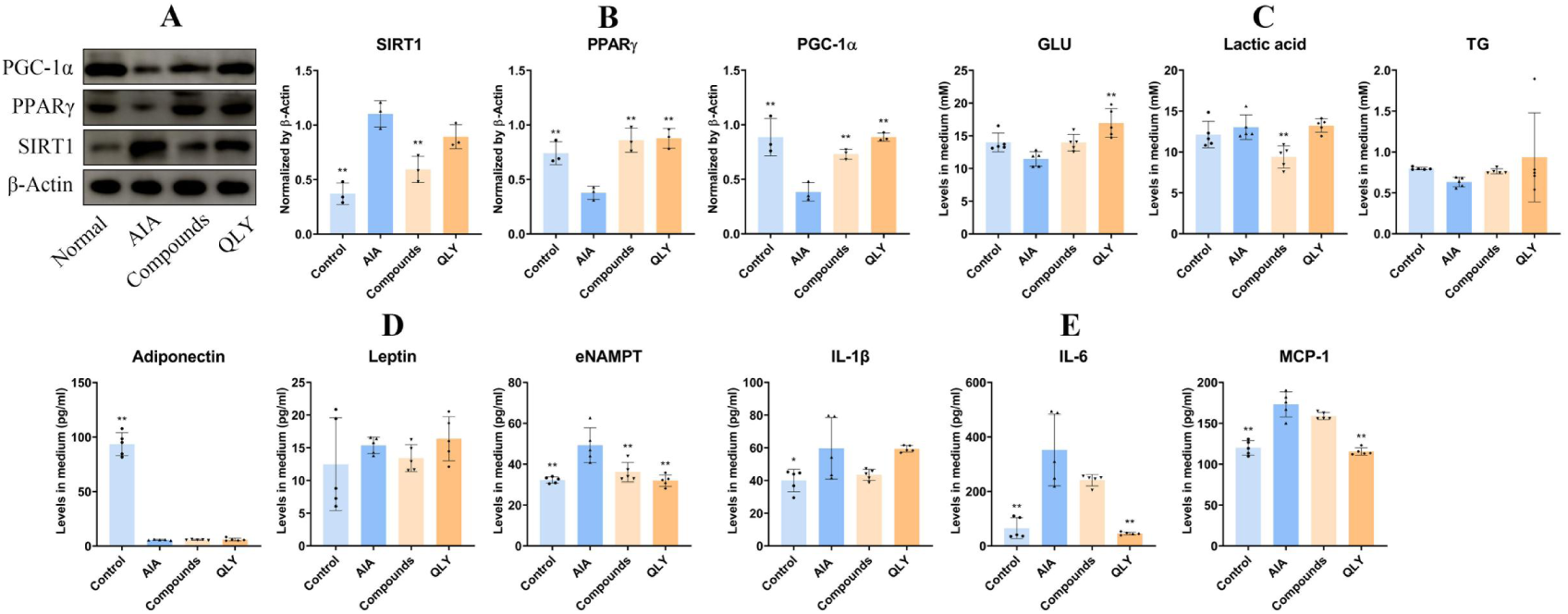
QLY-related SIRT1 inhibitors exerted the similar effects on pre-adipocytes to QLY-containing serum. A, WB assays performed using the pre-adipocytes cultured in vitro (they were first cultured by healthy, AIA model rats’ serum and then treated by the compounds or QLY-containing serum); B, the quantification results of assay A; C, levels of some key metabolites (GLU, TG and lactic acid) in the medium above; D, levels of the representative adipokines (leptin, adiponectin, and eNAMPT) in the medium; E, levels of the representative cytokines (IL-1β, IL-6 and MCP-1) in the medium. Statistical significance: *p < 0.05 and **p < 0.01 compared with AIA group.

Then, we intended to further confirm the beneficial consequences from SIRT1 regulation in the above experiments by regulating this signal. In WB experiments, SIRT1 overexpression perfectly mimicked AIA serum stimulation-caused changes in pre-adipocytes. The two treatments promoted expression of SIRT1 and tNAMPT proteins, and decreased expression of PPARγ and PGC-1α. Meanwhile, QLY-related compounds significantly reversed this expression change trend, and this outcome was partially offset by SIRT1 overexpression (**Figure 7A**). The compounds restored levels decline of extracellular GLU and TG in AIA serum-primed cells. Although SIRT1 overexpression didn’t significantly affect the nutrients, it impaired the effects of the compounds. It suggests that AIA condition had already sufficiently activated SIRT1, and QLY-induced SIRT1 inhibition curb energy utilization. Interestingly, we observed that SIRT1 overexpression and the chemical treatment synergistically inhibited lactic acid production (**Figure 7B**). It suggests that SIRT1 inhibition didn’t contribute to the down-regulation of glycolysis during QLY treatments, and there exists some other mechanisms mediating this outcome. Then, we tested some secreted factors sensitive to QLY treatment. SIRT1 overexpression can significantly reduce the expression of IL-6, indicating the anti-inflammatory role of SIRT1. But it didn’t affect MCP-1 and eNAMPT. We can infer that their production would be either unaffected by SIRT1 or promoted by it. QLY-related compounds reduced the secretion of IL-6 and MCP-1 a lot, but the effects were antagonized by SIRT1 overexpression more or less (**Figure 7C**).

**Figure 7.**
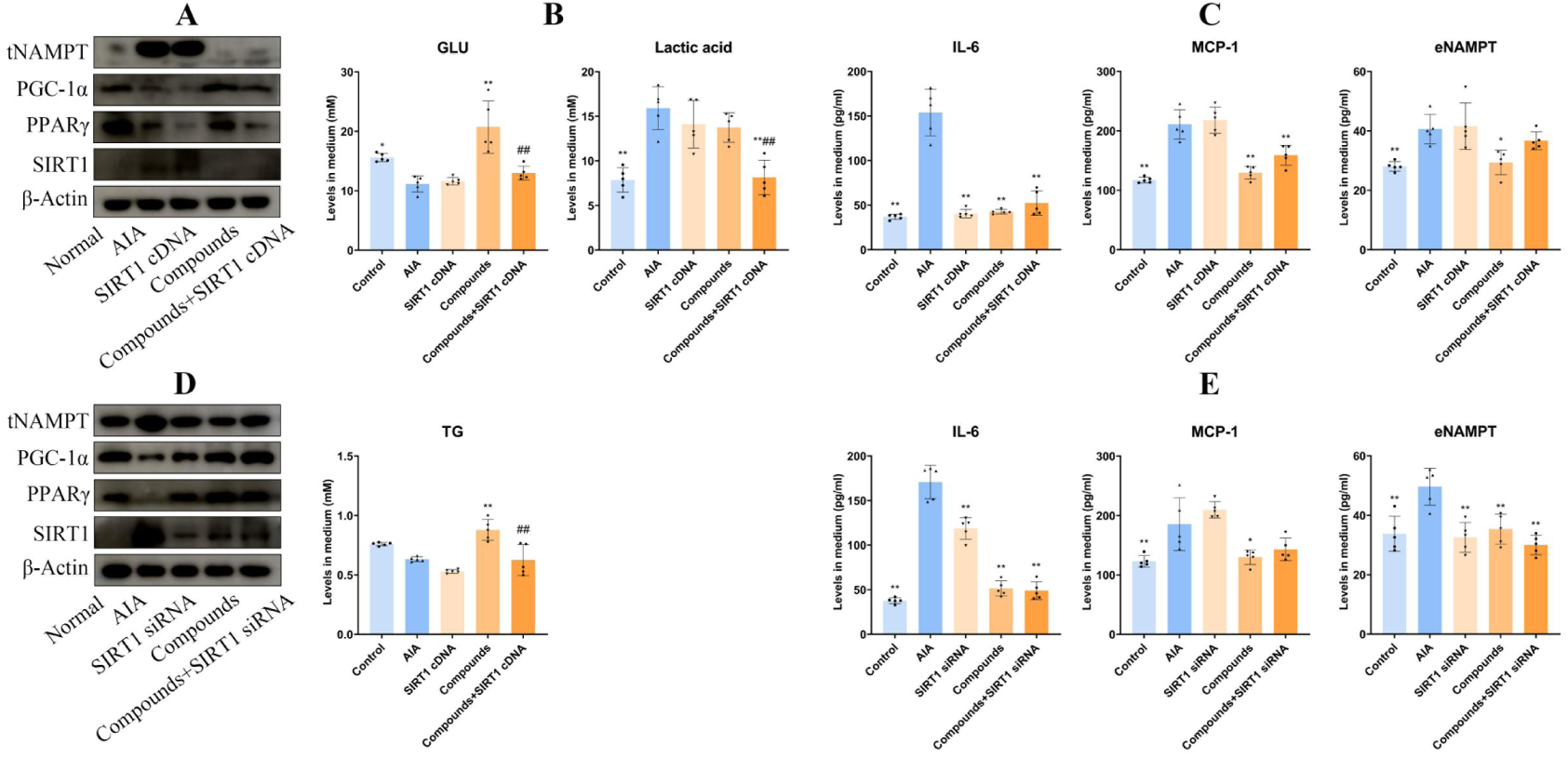
QLY-induced eNAMPT decrease in pre-adipocytes was attributed to SIRT1 inhibition. A, WB assays performed using some pre-adipocytes cultured in vitro (SIRT1 in some of them were overexpressed, and they were exposed to QLY-related SIRT1 inhibitors after different serums culture); B, levels of some key metabolites (GLU, TG and lactic acid) in the medium from assay A; C, levels of representative WAT-released substances (eNAMPT, IL-6 and MCP-1) in the medium; D, WB assays performed using some pre-adipocytes cultured in vitro (SIRT1 in some of them were silenced, and they were exposed to QLY-related SIRT1 inhibitors after different serums culture); E, levels of representative WAT-released substances in the medium from assay D. Statistical significance: *p < 0.05 and **p < 0.01 compared with AIA group; ^##^p < 0.01 compared with compounds group.

Obviously, SIRT1-silencing in the pre-adipocytes achieved an opposite outcome to SIRT1 overexpression. In fact, its effects were very similar to those induced by the compounds treatment (**Figure 7D**). Interestingly, SIRT1-silencing also reduced IL-6 production in the AIA serum-cultured pre-adipocytes, and it didn’t affect the effects of the compounds in this aspect. It demonstrates the complex facets of SIRT1 in immune regulation. Similar to SIRT1 overexpression, its silencing had no effect on MCP-1 either. This finding confirms that SIRT1 doses control MCP-1 expression. Importantly, these results hint that QLY suppressed the release of inflammatory cytokines in WAT possibly a way other than SIRT1 inhibition. As expected, SIRT1-silencing achieved a similar inhibitory effect on eNAMPT levels to QLY-related compounds (**Figure 7E**). It confirms that eNAMPT decrease is the main beneficial outcome from QLY-induced SIRT1 inhibition in WAT.

## Discussion

The autoimmune conditions of RA trigger persistent inflammation and tissue injuries. It makes immunosuppressive therapy aiming at synovitis a priority option for RA patients. The first-line drugs of RA therapies including corticosteroids, non-steroidal anti-inflammatory drugs, and disease-modifying anti-rheumatic drugs all work via this way.^40^ Accordingly, increasing pathological immune cells have been recognized, such as T/B lymphocytes, monocytes/macrophages, granulocytes, dendritic cells, and so on.^41^ However, none of immune regulation-based therapies can cure RA. In fact, many patients don’t respond well to them. Hence, there exist some other players in RA pathology.^46^ It is already known now that immune activation requires metabolism reprogramming. Hence, metabolic intervention would affect the prognosis of many diseases. This recognition highlights the role of metabolic organs.

RA patients are prone to develop metabolic disorders. Approximately 30% of RA patients exhibit central obesity.^43^ Because WAT is the main source of inflammatory cytokines, this phenomenon bridges RA with adipocyte dysfunction. We have recently confirmed that WAT is an amplifier of RA-related immune abnormalities, and showed that changes of certain pathways involved in adipocyte development and maturation have profound clinical implications.^44^ SIRT1/PPARγ then becomes a RA-related hotspot in this circumstance, because it decides both immune and immune functions of many tissues, including WAT. Our early studies showed that PPARγ activation was involved in the anti-rheumatic actions of QLY.^14,40^ Furthermore, we recently discovered that QLY contains some compounds that effectively inhibit SIRT1 activity.^29^ As well-known, PPARγ negatively regulates inflammation. We can anticipate the favorable consequences from QLY-induced PPARγ activation. However, how SIRT1 inhibition contributes to the therapeutic effects of QLY on RA is very difficult to be understood, as SIRT1 is usually recognized as a negative regulator of inflammation too. The current study provided some useful clues to elucidate this mystery from three aspects: (ⅰ) SIRT1 signaling status varies a lot in different organs/cells of rheumatic subjects, and these differed preconditions decide the different outcomes from SIRT1 regulation; (ⅱ) the reciprocal inhibition between PPARγ and SIRT1 promises a conditionally pro-inflammatory role of SIRT1; (ⅲ) not all the secreted substances are sensitive to SIRT1 changes in WAT, and QLY-induced SIRT1 inhibition improves immune milieu mainly by reducing eNAMPT production.

QLY is used to treat hot syndromes-associated RA subtype, which is mainly characterized by evident inflammation manifestations. Hence, we first studied crucial pathological changes in the inflammatory stages, and found inflammatory polarization of monocytes is a hallmark.^20^ Now available knowledge suggests that these cells drive RA progress basically in two ways: (ⅰ) they infiltrate into synovial tissues, where they produce excessive inflammatory cytokines and promote Th17 cell differentiation; (ⅱ) they readily differentiate into osteoclasts, and therefore participate in bone damage.^6^ It is reasonable to assume that QLY would inhibit inflammatory monocytes, and indeed we observed this anticipated outcome.^10,14^ Insufficient SIRT1 expression and activation are revealed to a cause of the uncontrolled inflammatory polarization of monocytes in rheumatic subjects.^23^ It is contradicted to QLY-held inhibitory effects on SIRT1. In fact, the direct effects of QLY on monocytes are weak.^10^ Our latest work confirms that the limited ingredients permeated into blood after oral QLY doses are unable change the immune phenotype of monocytes in AIA rats (unpublished). In this study, we also observed that QLY didn’t change the phenotype of circulating monocytes in AIA rats during remission stages (**Figure 1**). From this perspective, SIRT1 inhibitors in QLY won’t exacerbate inflammation in monocytes either. They preferentially impact metabolic organs like liver and WAT, where SIRT1 is usually up-regulated.^29^ It promises a possibility that QLY indirectly improves immune conditions of rheumatic subjects by inhibiting SIRT1 in WAT.

There are many evidences supporting the anti-rheumatic potentials of SIRT1. It inhibits the transcriptional activity of NF-κB and AP-1 mainly via deacetylation of the key subunits.^45^ Furthermore, SIRT1 favors oxidative phosphorylation but inhibit glycolysis, just as we observed in **Figure 7B**. Its activation therefore will undermine the metabolic reprogramming required by inflammatory monocytes.^24^ As the result, SIRT1 agonists demonstrate promising anti-RA effects.^23,46^ But we cannot ignore the possible negative role of SIRT1 in RA. Increased SIRT1 expression is observed in the joints of RA patients. Synovioblasts with elevated SIRT1 levels exhibit resistance to apoptosis and display particularly notable pathological activities.^19^ Additionally, SIRT1 promotes the differentiation of notorious Th17 cells.^47^ In WAT, SIRT1 may deteriorate immune milieu of rheumatic subjects by two possible ways: inhibiting PPARγ and promoting eNAMPT release.^18,22^ We confirm SIRT1 suppresses the expression of PPARγ in **Figure 7A**. Together with the evidences shown in **Figure 4A**, we conclude that SIRT1 over-activation in AIA rats directly accounts for the impaired expression of PPARγ in WAT. Consequently, the production of IL-1β and some other inflammatory cytokines was increased (**Figure 5B**). This theory successfully explains the tricky impacts of SIRT1 regulation on IL-6 production. SIRT1 overexpression will reduce IL-6 production because of the anti-inflammatory property of SIRT1. On the other side, SIRT1-silencing also causes IL-6 production decrease due to the increased expression of PPARγ (**Figure 6**). The clues also demonstrate the sophisticated impacts of SIRT1 regulation on the secretion profile of WAT.

We usually tend to believe that effective therapies would bring levels of all the pathological factors down. But this result is not guaranteed because their synthesis and release are controlled by different mechanisms. Cytokines production is typically regulated by immune pathways, and therefore SIRT1 affects their levels in the indirect manner. Sometimes, this indirect mechanism is blunt. For instance, MCP-1 was barely affected by the varied status of SIRT1 (**Figure 6**). Although QLY-related compounds decreased its levels in AIA serum-primed pre-adipocytes, it cannot be attributed to SIRT1 inhibition. QLY-induced decrease of inflammatory cytokines must be mediated by some other targets. Different from cytokines, adipokines are usually recognized as metabolism-related hormones, and therefore their production is basically controlled by metabolic regulators like SIRT1. RA-related researches obtained rather conflicting data about the changes of adipokines. In fact, we observed the differed leptin changes in **Figure 3** and **6**. eNAMPT increase under rheumatic conditions is one of the solid conclusions from the previous studies.^16,29,31,44^ eNAMPT has the potential in initiating inflammation in immune cells by activating TLR4 and promoting glycolysis.^16^ These facts indicate that it would be a promising anti-rheumatic target. However, it is unlikely that any agents would substantially impair its expression, because this constitutively expressed enzyme is indispensable for living cells and its deletion is lethal. In conformity to this, QLY didn’t affect iNAMPT expression in pre-adipocytes (**Figure 5A**). The dramatic decrease in tNAMPT observed in **Figure 4** then can only be explained by the impaired processing of eNAMPT (**Figure 6D**). Interestingly, this step is controlled by SIRT1.^22^ Apparently, SIRT1 inhibitors in QLY contribute a lot to the decreased eNAMPT secretion (**Figure 7**). As SIRT1 overexpression possibly causes various harms to RA patients, if QLY-induced SIRT1 inhibition is involved in some other aspects of its ant-rheumatic actions is worthy of investigation.

## Conclusion

Although QLY therapy didn’t affect immune milieu of AIA rats during the remission stages, it profoundly changed the functions of liver and WAT by inhibiting SIRT1. All the effects are weakened by NMN, which activated SIRT1 by increasing NAD supply. QLY-induced SIRT1 inhibition in WAT affected its secretion profile, achieving the significant decrease of eNAMPT levels. This change relieved inflammation in AIA rats. These findings were confirmed in experiments in vivo by treating pre-adipocytes with QLY-derived. In short, the current study shows that SIRT1 inhibitors in QLY reduce eNAMPT release and consequently ease RA via the interplay between immune cells and WAT.

## Abbreviation

RA: rheumatoid arthritis
WAT: white adipose tissues
eNAMPT: extracellular NAMPT
iNAMPT: intracellular NAMPT
tNAMPT: total NAMPT
TCM: Traditional Chinese Medicine
QLY: Qing-Luo-Yin
BCG: Bacillus Calmette-Guérin
IFA: incomplete Freund’s adjuvant
CFA: complete Freund’s adjuvant
GSH: reduced glutathione
MDA: malonaldehyde
SOD: superoxide dismutase
AKP: alkaline phosphatase
GSP: glycated serum protein
GLU: glucose
TG: triglyceride
T-CHO: total cholesterol
HDL-C: high-density lipoprotein cholesterol
LDL-C: low-density lipoprotein cholesterol
NMN: nicotinamide mononucleotide
WB: western-blot
IP: immunoprecipitation
RT-PCR: reverse transcription polymerase chain reaction
cDNA: complementary DNA
qPCR: quantitative PCR
BSA: bovine serum albumin

## Data Sharing Statement

All data generated or analyzed during this study are included in this article and its Supplementary Information.

## Ethics approval

The animal study was approved by the Ethical Committee of Wannan Medical College (WNMC-AWE-2023358). The use of animals was adhered to the National Institutes of Health Guide for the Care and Use of Laboratory animals (2011), and all the experiment methods were in accordance with ARRIVE guidelines.

## Author contributions

Jian Zuo and Kui Yang conceived the idea; Dan-Dan Wang and Meng-Ke Song performed majority of the experiments; Qin Yin and Wen-Gang Chen assisted all the experiments; Jian Zuo wrote the manuscript; Opeyemi Joshua Olatunji proof read the manuscript. All the authors gave final approval of the version to be published.

